# NLRP11 is required for canonical NLRP3 and non-canonical inflammasome activation during human macrophage infection with mycobacteria

**DOI:** 10.1101/2024.12.11.627830

**Authors:** Mateusz Szczerba, Akshaya Ganesh, María Luisa Gil-Marqués, Volker Briken, Marcia B. Goldberg

**Affiliations:** Center for Bacterial Pathogenesis, Division of Infectious Diseases, Department of Medicine, Massachusetts General Hospital, Boston, MA, USA; Department of Microbiology, Blavatnik Institute, Harvard Medical School, Boston, MA, USA; Department of Cell Biology and Molecular Genetics, University of Maryland, College Park, MD, USA; Broad Institute of MIT and Harvard, Cambridge, MA, USA; Department of Immunology and Infectious Diseases, Harvard T. H. Chan School of Public Health, Boston, MA, USA

**Keywords:** Mycobacteria, innate immunity, macrophages, inflammasomes, NLRP11, NLRP3

## Abstract

The NLRP11 protein is only expressed in primates and participates in the activation of the canonical NLRP3 and non-canonical NLRP3 inflammasome activation after infection with gram-negative bacteria. Here, we generated a series of defined NLRP11 deletion mutants to further analyze the role of NLRP11 in NLRP3 inflammasome activation. Like the complete NLRP11 deletion mutant (*NLRP11^-/-^*), the NLRP11 mutant lacking the NACHT and LRR domains (*NLRP11*^Δ*N_LRR*^) showed reduced activation of the canonical NLRP3 inflammasome, whereas a pyrin domain mutant (*NLRP11*^Δ*PYD*^) had no effect on NLRP3 activation. The *NLRP11^-/-^* and *NLRP11*^Δ*N_LRR*^ mutants but not the *NLRP11*^Δ*PYD*^ mutant also displayed reduced activation of caspase-4 during infection with the intracytosolic, gram-negative pathogen *Shigella flexneri*. We found that the human adapted, acid-fast pathogen *Mycobacterium tuberculosis* and the opportunistic pathogen *M. kansasii* both activate the non-canonical NLRP11 inflammasome in a caspase-4/5-dependent pathway. In conclusion, we show that NLRP11 functions in the non-canonical caspase-4/5 inflammasome activation pathway and the canonical NRLP3 inflammasome pathway, and that NLRP11 is required for full recognition of mycobacteria by each of these pathways. Our work extends the spectrum of bacterial pathogen recognition by the non-canonical NLRP11-caspase4/5 pathway beyond gram-negative bacteria.

**IMPORTANCE:** The activation of inflammasome complexes plays a crucial role in intracellular pathogen detection. NLRP11 and caspase-4 are essential for recognizing lipopolysaccharide (LPS), a molecule found in gram-negative bacteria such as the human pathogens *Shigella* spp., which activate both canonical NLRP3 and non-canonical inflammasome pathways. Through a series of deletion mutants, we demonstrate that the NACHT and LRR domains of NLRP11, but not its pyrin domain, are critical for detection of *S. flexneri*. Notably, our research reveals that the acid-fast bacterium *M. tuberculosis* is also detected by NLRP11 and caspase-4, despite not producing LPS. These findings significantly expand the range of pathogens recognized by NLRP11 and caspase-4 to now include acid - fast bacteria that do not contain LPS and underscore the versatility of these innate immune components in pathogen detection.

## INTRODUCTION

Intracellular pathogens are detected by host cell cytosolic Nucleotide-binding Oligomerization Domain and Leucine-rich Repeat-containing receptors (NLRs) or Absent in Melanoma 2-like receptors (ALRs), which trigger the activation of an inflammasome complex (1–5). In addition to the respective NLR or ALR, this complex typically comprises an adaptor component (ASC) and pro-caspase-1.

Upon full activation, the inflammasome complex contains active caspase-1, which cleaves pro-IL-1β to generate mature IL-1β. Caspase-1 also cleaves gasdermin D (GSDMD), releasing an N-terminal (NT-GSDMD; GSDMD p30) fragment that forms pores in the cell membrane through which IL-1β exits the cell (6–10). The process culminates with the activation of NINJ1, a membrane disrupting protein that leads to plasma membrane rupture (11–13). This final stage marks the completion of pyroptosis, a specific and inflammatory form of cell death.

Activation of the canonical NLRP3 inflammasome is triggered by various intracellular stressors, including potassium efflux, which occurs with GSDMD pore formation or when cells are exposed to certain bacterial toxins that damage the cell membrane, or increased reactive oxygen species (ROS), a common cellular response to intracellular bacteria (14–21). The non-canonical inflammasome pathway relies on the recognition of lipopolysaccharide (LPS) by inflammatory pro-caspase-4/5 (humans) or their murine homologue pro-caspase- 11 (22–25). LPS binding leads to caspase activation and subsequent cleavage of GSDMD. The resulting NT-GSDMD pores facilitate K^+^ efflux, which in turn triggers NLRP3 inflammasome activation. Humans but not mice express NLRP11, which is required for efficient activation of the non-canonical inflammasome in response to intracellular LPS (26), as well as canonical NLRP3 inflammasome activation (27). In the current study, we determined the importance of the NLRP11 pyrin, NACHT, and/or LRR domains for their function in LPS recognition, IL-1β production and pyroptosis induction.

*Mycobacterium tuberculosis* is a human adapted, facultative intracellular pathogen that activates the NLRP3 inflammasome of infected macrophages (28). IL-1β is of great importance for a protective host response to *M. tuberculosis* infections (28, 29). *M. tuberculosis* evolved to evade recognition by host cell inflammasomes (28). The *M. tuberculosis* phospholipid phosphatase (PtpB) cleaves cell membrane phosphatidylinositol-4 phosphate which reduces insertion of NT-GSDMD into the plasma membrane and this limits pyroptosis and IL-1β secretion (30). The *M. tuberculosis* protein kinase (PknF) limits activation of the NLRP3 inflammasome via an unknown mechanism (31). The Rv3364c protein suppresses host cathepsin G activation, leading to reduced inflammasome activation (32). Despite all these inhibitory mechanisms, the host macrophage is still able to sense intracellular *M. tuberculosis* via activation of the NLRP3 inflammasome. *M. tuberculosis* is an acid-fast bacterium that does not express LPS but is atypical because it has an outer membrane composed of mycobacterial lipids and lipoproteins (33). Since *M. tuberculosis* is a human adapted pathogen and NLRP11 is only expressed in primates, we hypothesized the NLRP11 might participate in the recognition of mycobacteria.

## RESULTS

### NLRP11 is required for activation of the canonical NLRP3 inflammasome in human macrophages

We previously found that cells lacking the coding sequence of the NACHT and LRR domains of NLRP11 display defects in activation of the non-canonical inflammasome in response to cytosolic LPS (26). To further evaluate the contributions of NLRP11 to innate immune signaling, we generated a series of defined domain deletions in NLRP11 in human THP-1 monocytes (Fig. 1A): using CRISPR-Cas9 technology, we deleted the *NLRP11* pyrin domain [PYD] (*NLRP11*^Δ*PYD*^), the NACHT and leucine-rich repeat [LRR] domains (*NLRP11*^Δ*N_LRR*^), or the entire *NLRP11* coding sequence (*NLRP11^-/-^*) (Fig. S1 in the supplemental material). We generated these mutations using Cas9 ribonucleoproteins, which are associated with long-term genome stability, isolated clonal derivatives by limiting dilution and carried forward only those clonal lineages that contained a homozygous out-of-frame deletion.

**FIG 1.**
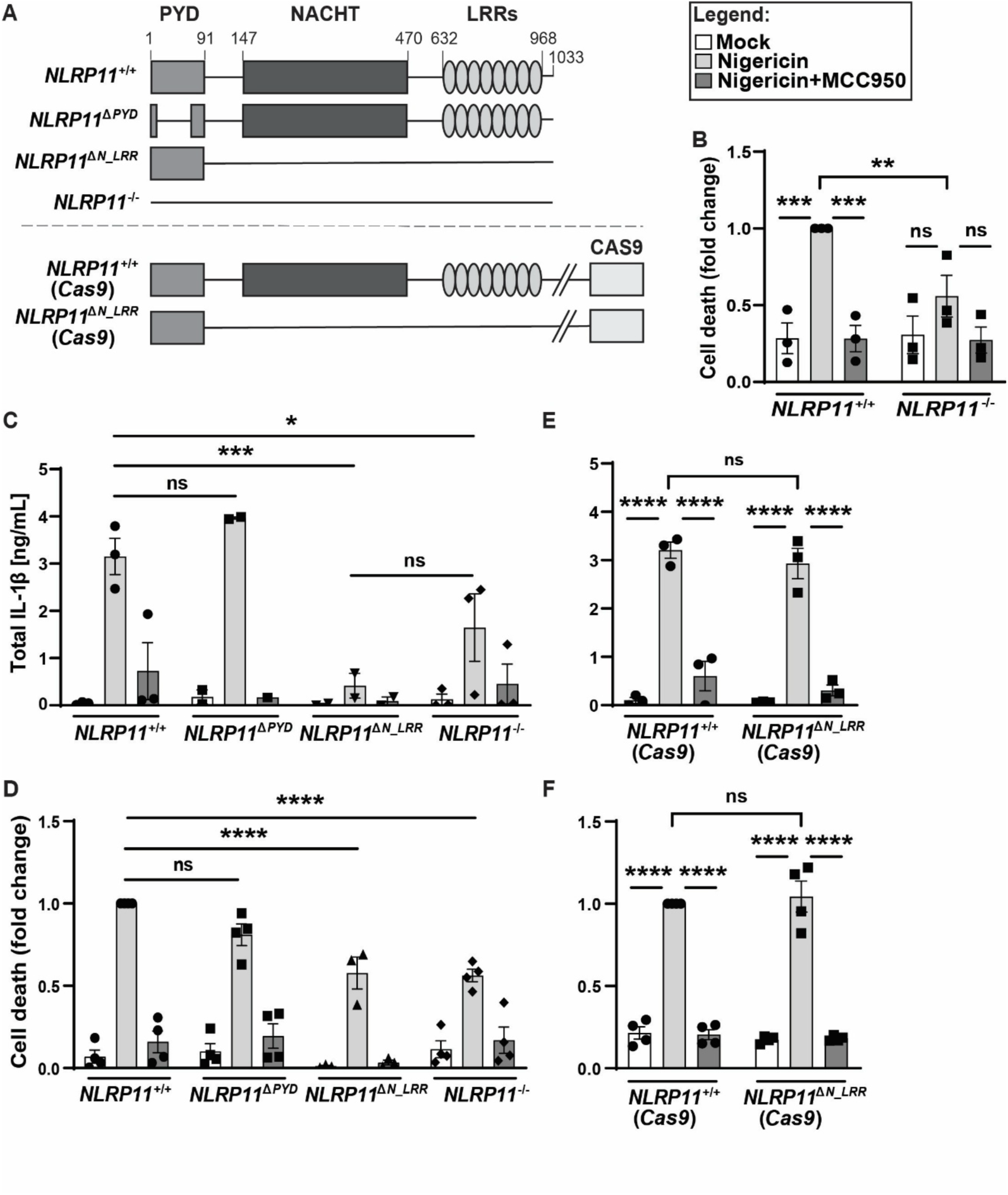
Efficient activation of the canonical NLRP3 inflammasome depends on the NLRP11 NACHT and/or LRR domains. (A) Schematic of *NLRP11* mutants generated in THP-1 macrophages. (B) Activation of cell death in *NLRP11*^+/+^ and *NLRP11*^-/-^ macrophages by the NLRP3 agonist nigericin. Cell death, measured via LDH release assay at 90 min, is shown normalized to nigericin-treated *NLRP11*^+/+^ macrophages. The level of cell death in *NLRP11*^+/+^ macrophages treated with nigericin approximated 30%. Where indicated, cells were pretreated with the NLRP3 inhibitor MCC950. (C) Levels of IL-1β released into the cell culture media at 90 min of nigericin treatment with or without addition of MCC950. (D) Cell death, as in (B). (E-F) Levels of released IL-1β (E), as in (C), and cell death (F), as in (B), in macrophages with stably integrated Cas9. ns, not significant; *, *P* < 0.05; **, *P* < 0.01; ***, *P* < 0.001; ****, *P* < 0.0001; by two-way ANOVA. Shown are independent biological replicates. Error bars represent the standard error of the mean. PYD, pyrin domain; NACHT, NAIP, C2TA, HET-E and TP1 domain; LRR, leucin-rich repeat domain.

We assessed the impact of deletions of *NLRP11* on activation of the canonical NLRP3 inflammasome. In ribonucleoprotein-generated *NLRP11^-/-^* macrophages, although NLRP3 levels were not reduced (Fig. S1), cell death upon treatment with the NLRP3 agonist nigericin was significantly decreased compared to nigericin-treated WT macrophages (Fig. 1B). Activation of caspase-1, which involves cleavage and release of its active domains, and activation of GSDMD, which involves its cleavage, were significantly reduced in *NLRP11^-/-^* macrophages (Fig. S2). In parallel, release of IL-1β, a pro-inflammatory cytokine effector of inflammasome activation, was reduced in *NLRP11^-/-^* macrophages (Fig 1C).

To test whether these phenotypes were specifically due to activation of NLRP3, we treated cells with the NLRP3 inhibitor MCC950. Treatment with MCC950 rescued nigericin-induced cell death, processing of caspase-1 and GSDMD, and release of IL-1β and IL-18 to the levels of mock-treated THP-1s (Fig. 1B-C, and Fig. S2), indicating that the observed phenotypes were due to activation of NLRP3.

### NACHT and/or LRR domains are required for NLRP11-dependent activation of the NLRP3 inflammasome

To determine the impact of specific domains of NLRP11 on NLRP3 signaling, we assessed activation of cell death and cytokine release in ribonucleoprotein-generated *NLRP11*^Δ*PYD*^ and *NLRP11*^Δ*N_LRR*^ macrophages. Deletion of NACHT and LRR domains, but not the PYD domain alone, was associated with reduced nigericin-induced cell death (Fig. 1D) and reduced release of IL-1β (Fig. 1C). In addition, in *NLRP11*^Δ*N_LRR*^ macrophages, processing of caspase-1 and GSDMD was significantly reduced (Fig. S2). In contrast, in *NLRP11*^Δ*PYD*^ macrophages, caspase-1 was activated at levels similar to those observed in *NLRP11^+/+^* macrophages, and activation of GSDMD was only mildly decreased. These findings indicate that the NLRP11 NACHT and/or LRR domains are required for efficient NLRP3 inflammasome activation, consistent with previous data demonstrating a requirement for the NLRP11 NACHT and LRR domains for oligomerization of NLRP3 (27). The findings also suggest that the involvement of NLRP11 in NLRP3 activation does not strictly depend on the NLRP11 pyrin domain.

As we found previously (26), nigericin activation of NLRP3 was not impacted in *Cas9*-integrated *NLRP11*^Δ*N_LRR*^ macrophages (*NLRP11*^Δ*N_LRR*^ (*Cas9*), Fig. 1E-F). This contrasts with what we observe in *NLRP11*Δ*^N_LRR^* macrophages that lack *Cas9* (*NLRP11*^Δ*N_LRR*^, Fig. 1C-D). This result suggests that, as relates specifically to NLRP3 signaling, the observed lack of impact of NLRP11 NACHT and LRR disruption on NLRP3 in the *Cas9*-integrated cell lineage is not due to the maintenance of the NLRP11 pyrin domain; rather, the lack of impact on NLRP3 in this lineage likely results from non-NLRP11 genetics and suggests that genome-level changes resulting from the chronic presence of *Cas9* contributed to this phenotype.

### Activation of the NLRP3 inflammasome by infection with the acid-fast bacterium *M. tuberculosis* depends on NLRP11

Whether NLRP11 is involved in recognition of acid-fast or other gram-positive bacteria and mounting of a cellular inflammatory response to them has not been studied. IL-1β produced by THP-1 macrophages infected with *M. tuberculosis* is dependent on NLRP3 (34, 35). NLRP11 is also recruited by NLRP3 in the process of its activation (27).

Therefore, we hypothesized that in *NLRP11*^-/-^ THP-1 macrophages infected by *M. tuberculosis*, IL-1β production might be lower than in WT macrophages. To test this hypothesis, we determined levels of IL-1β release during infection with *M. tuberculosis* of macrophages containing or lacking NLRP11. As a readout for inflammasome activation during *M. tuberculosis* infection, cell death is less useful than IL-1β release because few *M. tuberculosis*-infected macrophages die. At 8 and 24 hours of infection, significantly less IL-1β was released from *NLRP11*^-/-^ macrophages than from WT macrophages (Fig. 2A-B). To test whether the observed IL-1β release is dependent on activation of NLRP3, infected cells were treated with MCC950. *M. tuberculosis*-infected *NLRP11*^-/-^ macrophages treated with MCC950 display a decrease of IL-1β compared with those not treated with MCC950 (Fig. 2A-B), indicating a requirement for NLRP3, which could be direct or indirect.

**FIG 2.**
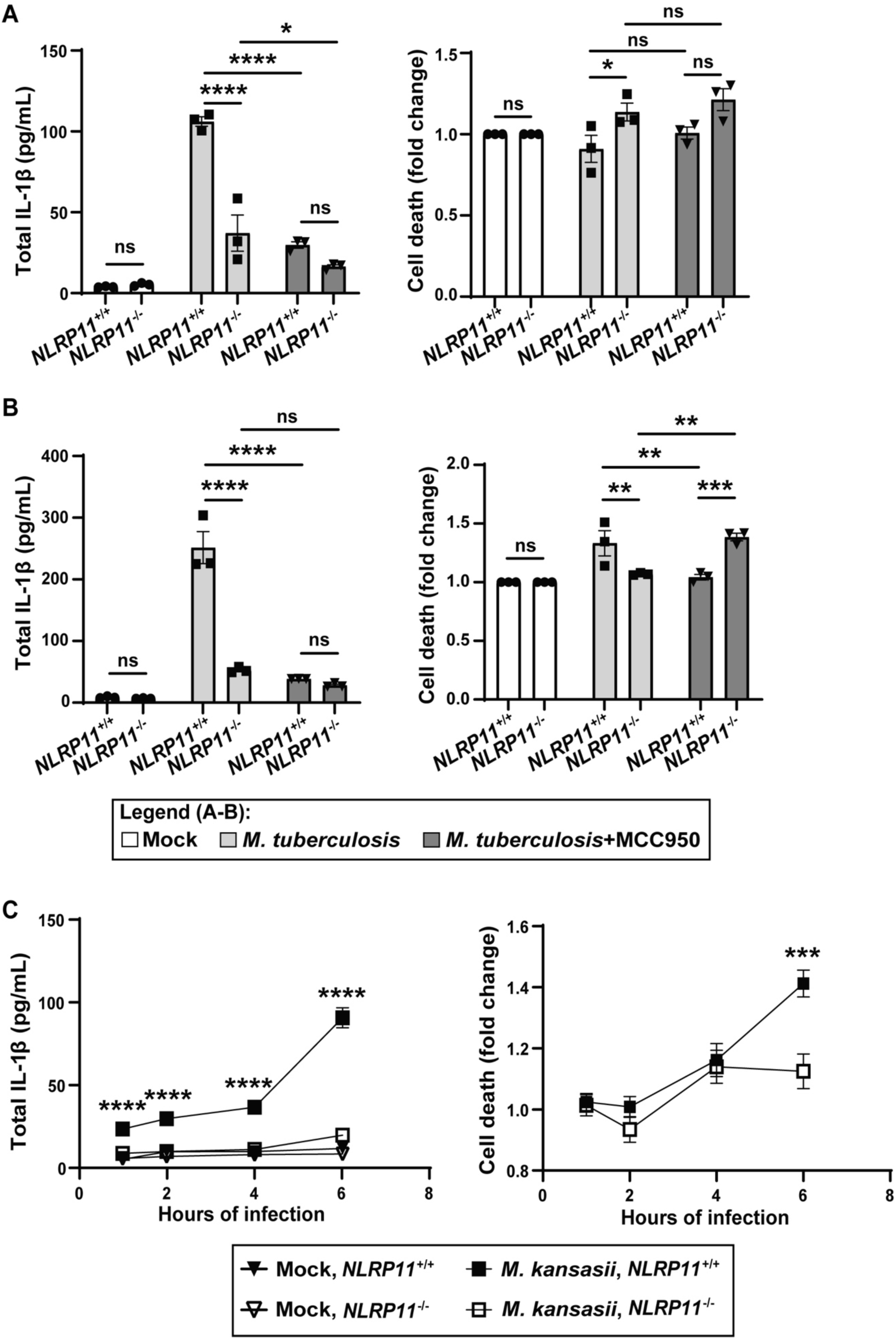
NLRP3 activation by *M. tuberculosis* and *M. kansasi* in THP-1 macrophages is dependent on NLRP11. (A and B) IL-1β release and cell death of *NLRP11*^-/-^ and WT THP-1 macrophages at 8 h (A) or 24 h (B) of infection with *M. tuberculosis* (MOI 10) with or without addition of MCC950. (C) IL-1β release and cell death of *NLRP11*^-/-^ and WT THP-1 macrophages over 1-6 h of infection with *M. kansasii*. Cell death was normalized to cell lineage-specific uninfected group. ns, not significant; *, *P* < 0.05; **, *P* < 0.01; ***, *P* < 0.001; ****, *P* < 0.0001; by two-way ANOVA (A-B) or paired parametric t-tests (C). Shown are independent biological replicates. Error bars represent the standard error of mean.

As for *M. tuberculosis*, IL-1β produced by THP-1 macrophages infected with *M. kansasii*, an opportunistic pathogen that is a close relative of *M. tuberculosis*, is dependent on NLRP3 (36). Release of IL-1β during infection with *M. kansasii* was dependent on NLRP11 (Fig. 2C, left graph), similar to the phenotype observed for *M. tuberculosis*-infected cells. At late times of *M. kansasii* infection, cell death was also partially dependent on NLRP11 (Fig. 2C, right graph).

Whereas IL-1β release from *NLRP11*^-/-^ macrophages was significantly reduced at 1, 2, 4 and 6 h of infection, cell death was significantly lower only at 6 h of infection (Fig. 2C).

### NLRP11 NACHT and/or LRR domains are required for activation of the non-canonical inflammasome by the gram-negative bacterium *Shigella flexneri*

We previously found that NLRP11 is required for activation of the non-canonical inflammasome during infection with a variety of gram-negative bacteria, including *S. flexneri* (26). Cell death during infection with *S. flexneri* was unaffected by deletion of the pyrin domain (PYD), whereas deletion of the NACHT and LRR domains or deletion of the entire *NLRP11* locus resulted in 40-50% decreases in macrophage cell death (Fig. 3A). Furthermore, release of pro-inflammatory cytokines IL-1β and IL-18 was reduced up to 10-fold in macrophages lacking the NACHT and LRR domains (*NLRP11*^Δ*N_LRR*^) or lacking all of NLRP11 (*NLRP11^-/-^*), but not in macrophages lacking only the PYD (*NLRP11*^Δ*PYD*^) (Fig. 3B).

**FIG 3.**
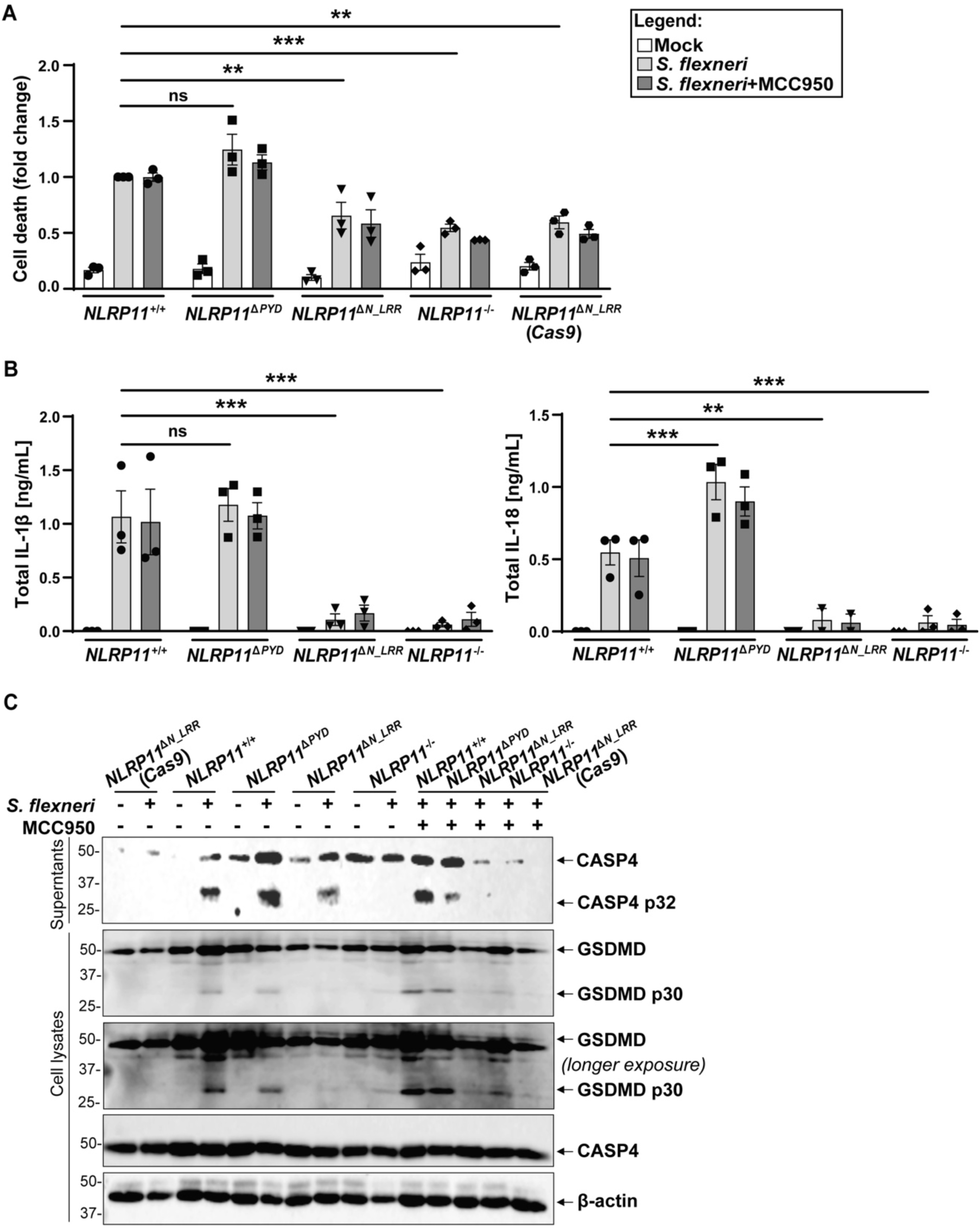
NACHT and/or LRR domains are required for NLRP11-dependent non-canonical inflammasome activation during *S. flexneri* infection. (A) Activation of cell death in indicated *NLRP11* mutant cell lines at 3 h of infection with *S. flexneri* (MOI 10). Cell death is shown normalized to *S. flexneri*-infected NLRP11^+/+^ macrophages. The level of cell death in infected *NLRP11*^+/+^ macrophages approximated 15-20%. Where indicated, cells were pretreated with the NLRP3 inhibitor MCC950. (B) Levels of IL-1β and IL-18 released into the cell culture media from (A). (C) Cleaved caspase-4 (CASP4 p32) released into the cell culture media and intracellular cleaved GSDMD (p30) under same conditions. Representative western blots. ns, not significant; **, P < 0.01; ***, P < 0.001; by two-way ANOVA. Shown are independent biological replicates (A-B). Error bars represent the standard error of the mean.

To test whether the observed defect in activation of pyroptosis in *NLRP11*^Δ*N_LRR*^ and *NLRP11^-/-^* macrophages resulted from a defect in activation of the non-canonical inflammasome, we examined processing of caspase-4 and GSDMD. Activation of caspase-4 was reduced approximately 2-fold in *NLRP11*^Δ*N_LRR*^ and *NLRP11^-/-^* macrophages but not reduced in *NLRP11*^Δ*PYD*^ macrophages (Fig. 3C and Fig. S3). Levels of full-length pro-caspase-4 in cell lysates were unaffected, indicating that the decrease in caspase-4 released from *NLRP11*^Δ*N_LRR*^ and *NLRP11^-/-^* macrophages was not due to changes in its expression or stability.

Similarly, processing of GSDMD was significantly reduced in *NLRP11*^Δ*N_LRR*^ and *NLRP11^-/-^* macrophages but not in *NLRP11*^Δ*PYD*^ macrophages, yet levels of full-length GSDMD were unaltered (Fig. 3C and Fig. S3). Thus, the NACHT and/or LRR domains are required for efficient activation of the NLRP11-dependent non-canonical inflammasome during infection with *S. flexneri*.

### Activation of the non-canonical inflammasome by *S. flexneri* is independent of NLRP3

To test whether the NLRP11-dependent activation of the non-canonical inflammasome is dependent on NLRP3, we examined inflammasome activation in *S. flexneri*-infected *NLRP11*^Δ*N_LRR*^ macrophages pre-treated with MCC950. Treatment with MCC950 did not impact the levels of cell death, cytokine release, or processing of caspase-4 and GSDMD (Fig. 3A-C, and Fig. S3). Therefore, NLRP3 is not required for activation of the NLRP11-dependent non-canonical inflammasome during *S. flexneri* infection.

### Activation of the non-canonical inflammasome by *M. tuberculosis* is dependent on NLRP11

The *NLRP11*^Δ*N_LRR*^ (*Cas9*) cell line accurately reports on the role of NLRP11 in non-canonical caspase-4 activity, while displaying normal activation of NLRP3 in response to nigericin (26). Therefore, to determine whether NLRP11 is required for non-canonical caspase-4 inflammasome activity during mycobacterial infection, we compared IL-1β release from mycobacteria-infected WT (*Cas9*) to that from mycobacteria-infected *NLRP11*^Δ*N_LRR*^ (*Cas9*) macrophages. *NLRP11*^Δ*N_LRR*^ cells infected with *M. tuberculosis* or *M. kansasii* showed significantly diminished IL-1β release at all time points (Fig. 4A-B), whereas those infected with the non-pathogenic *M. smegmatis* showed increases in IL-1β release over time that were similar to *M. kansasii* but which were not reduced in the *NLRP11*^Δ*N_LRR*^ cells at any time point (Fig. 4C). These data indicate that IL-1β released during *M. tuberculosis* and *M. kansasii* infection is dependent, at least partially, on the NACHT and/or LRR domains of NLRP11. Cell death was not significantly different between cell lines during infection with any of the three mycobacterial species.

**FIG 4.**
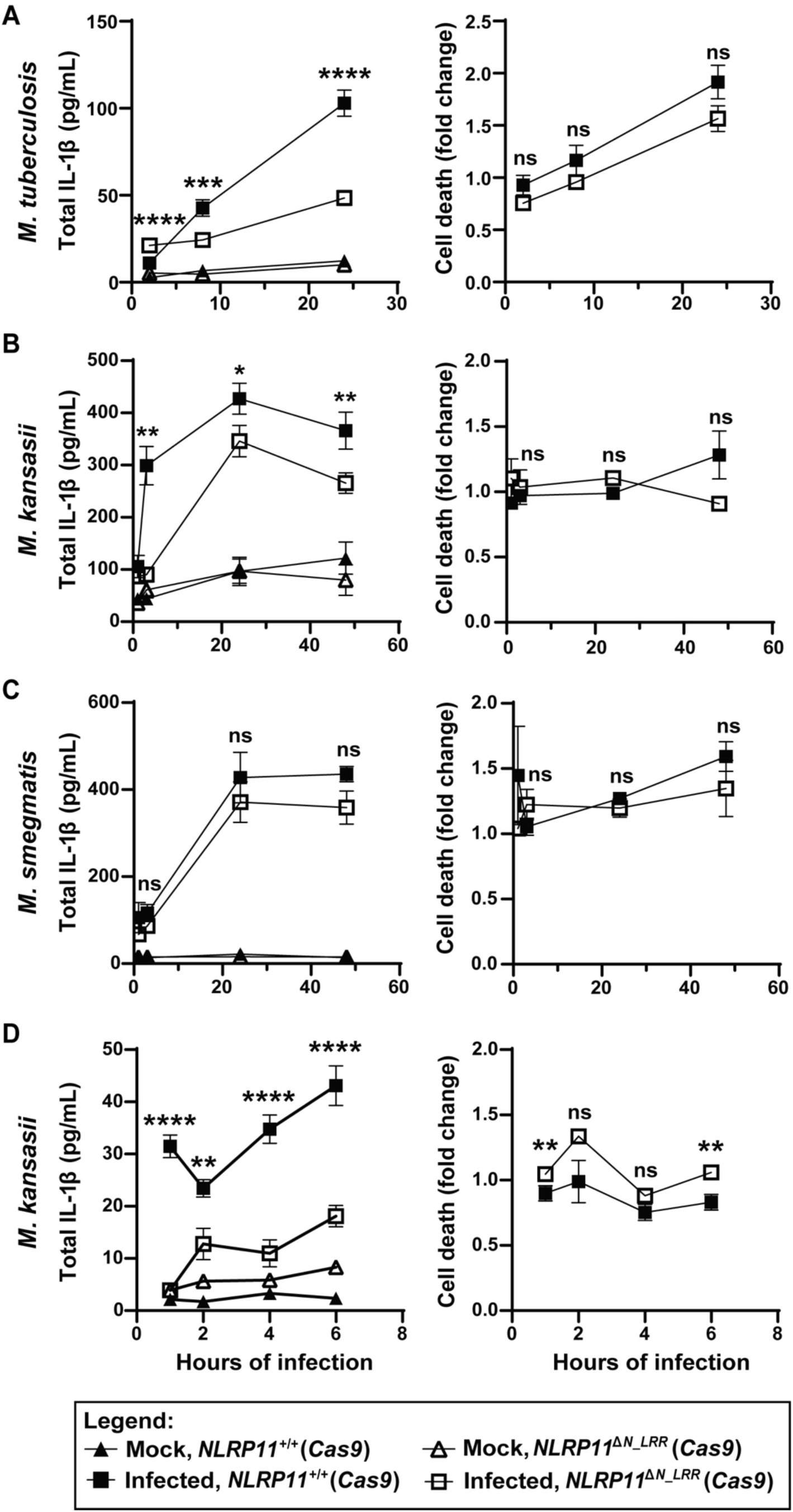
Mycobacteria activate the non-canonical inflammasome pathway. IL-1β release and cell death of *NLRP11*^+/+^ (*Cas9*) and *NLRP11*^ΔN_LRR^ (*Cas9*) mutant THP-1 macrophages infected with *M. tuberculosis* (A), *M. kansasii* (B, D), or *M. smegmatis* (C) (MOI 10), at the indicated timepoints. Cell death was normalized to cell lineage-specific uninfected group. The level of cell death in *NLRP11*^ΔN_LRR^ (*Cas9*) cells approximated 10-20%. ns, not significant; **, *P* < 0.01; ***, *P* < 0.001; ****, *P* < 0.0001; by paired parametric t-tests, comparing infected *NLRP11*^+/+^ (*Cas9*) to infected *NLRP11*^ΔN_LRR^ (*Cas9*). Shown are independent biological replicates. Error bars represent the standard error of mean.

To further examine the kinetics of NLRP11 dependent activation of the non-canonical inflammasome, we assayed IL-1β release in *M. kansasii*-infected WT and *NLRP11*^Δ*N_LRR*^ (*Cas9*) macrophages at 1, 2, 4 and 6 hours of infection. At every time point, significantly lower levels of IL-1β were released from *NLRP11*^Δ*N_LRR*^ macrophages than their WT counterparts (Fig. 4D), indicating that NLRP11 is required for IL-1β release even early during infection. As above (Fig. 4B), at early times, NLRP11-dependent differences in cell death are minimal (Fig. 4D).

The use of the NLRP11^ΔN_LRR^ (*Cas9*) mutant THP-1 cells is consistent with but not proof that *M. tuberculosis* and *M. kansasii* infection induces activation of the non-canonical inflammasome. In humans, the non-canonical inflammasome involves either or both of caspase-4 and caspase-5. During NLRP11-dependent activation of the non-canonical inflammasome, NLRP11 is required for downstream activation of at least caspase-4 (26); in this study, caspase-5 was not examined. To directly test whether mycobacteria activate the non-canonical inflammasome, we tested whether macrophages that lack both caspase-4 and caspase-5 support mycobacterium-induced IL-1β release and cell death. At 4 hours of infection with *M. kansasii* (Fig. 5A, left graph) and 24 hours of infection with *M. tuberculosis* (Fig. 5B, left graph), the absence of caspases-4 and -5 was associated with a complete absence of IL-1β release. Paralleling the impact of caspases-4 and -5 on IL-1β release, during infection with *M. kansasii*, their absence was associated with a two-fold reduction in cell death to baseline (Fig. 5A, right graph). During infection with *M. tuberculosis*, the impact of their absence on cell death was minimal (Fig. 5B, right graph). In WT THP-1 macrophages, specific to infection with *M. tuberculosis* was a small amount of caspase-4 processing and release that was not blocked by inhibition of NLRP3 with MCC950 (Fig. S4). These data indicate that IL-1β secretion during *M. tuberculosis* and *M. kansasii* infection is dependent on the non-canonical caspase-4/5 inflammasome.

**FIG 5.**
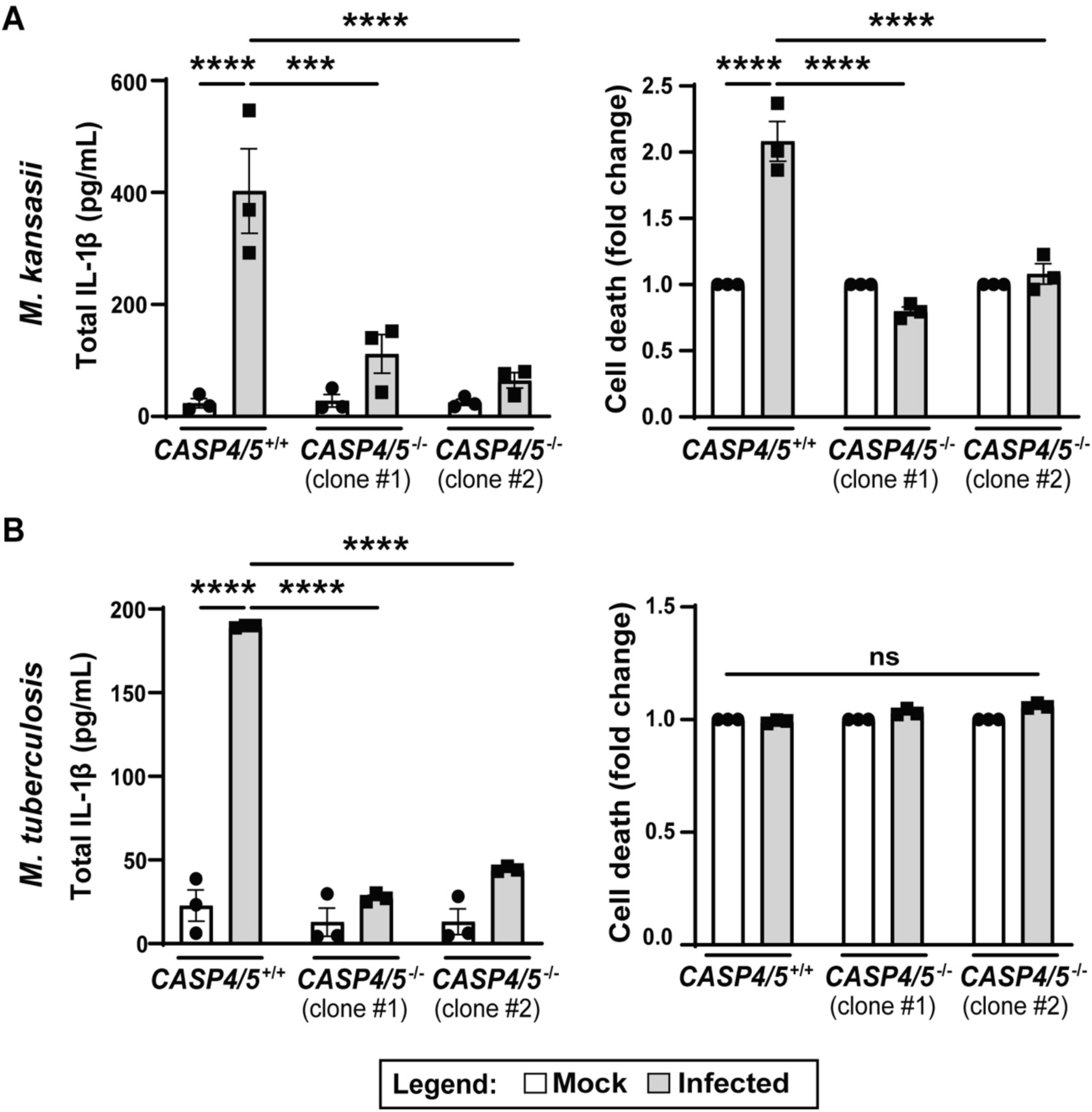
Caspase-4/5 are required for recognition of mycobacteria in human macrophages. IL-1β release and cell death of WT and two distinct clones of *caspase-4/5*^-/-^ (*CASP4/5*^-/-^) THP-1 macrophages infected with *M. kansasii* (A) or *M. tuberculosis* (B) (MOI 10). For *M. kansasii*, assays were performed at 4 h of infection (A), and for *M. tuberculosis* at 24 h of infection (B). Cell death in infected *CASP4/5*^-/-^ macrophages approximated 20-60%. ns, not significant; ***, *P* < 0.001; ****, *P* < 0.0001; by two-way ANOVA. Shown are independent biological replicates. Error bars represent the standard error of mean.

## DISCUSSION

Two recent studies establish that NLRP11 is required for efficient activation of the non-canonical caspase-4/5 inflammasome in macrophage-like THP-1 cells and primary human macrophages (26, 27). These studies demonstrated that NLRP11 is required for non-canonical inflammasome activation in response to cytosolic LPS (26, 27) and infection with gram-negative bacteria (26). Our findings herein extend our understanding of NLRP11 function by demonstrating that NLRP11 is required for efficient activation of the non-canonical inflammasome by the acid-fast bacteria *M. tuberculosis* and *M. kansasii* (Fig. 6).

**FIG 6.**
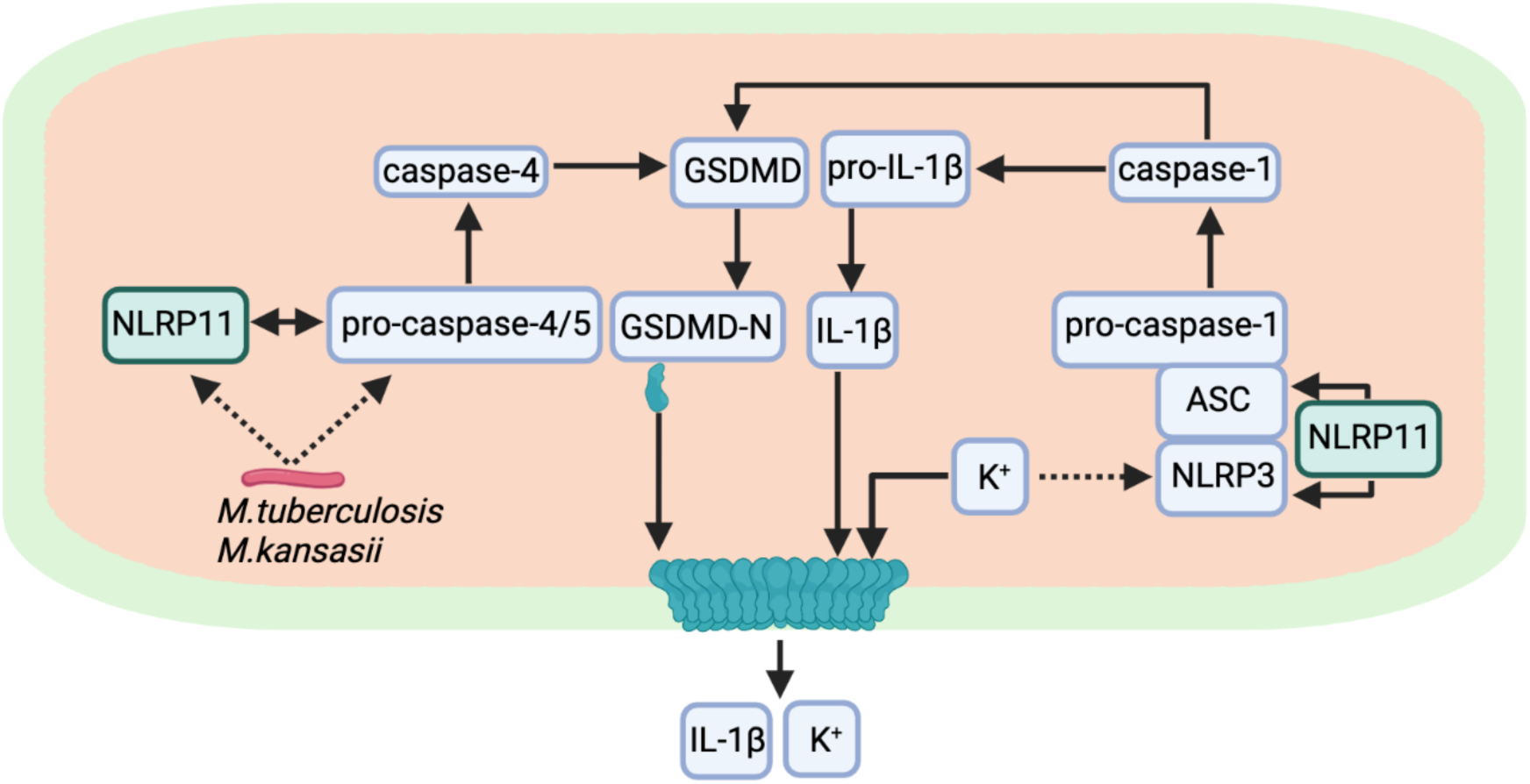
Role of NLRP11 in inflammasome activation during mycobacterial infection. NLRP11 displays dual function in activation of inflammasome responses in human macrophages infected with mycobacteria. NLRP11 and caspase-4 are required for efficient mycobacterial activation of the non-canonical inflammasome. Active caspase-4 cleaves GSDMD to release the N-terminal fragment (GSDMD-N) which forms pores in the plasma membrane. These pores lead to K^+^ efflux, a known trigger for activation of NLRP3. NLRP11 is required for efficient activation of the NLRP3 inflammasome, which leads to activation of caspase-1. Caspase-1 efficiently cleaves pro-IL-1β and GSDMD leading to more pore formation and release of mature IL-1β.

The present study offers clarification on previously conflicting findings regarding the role of NLRP11 role in activation of the canonical NLRP3 inflammasome: Gangopadhyay et al. reported that NLRP11 is required for NLRP3 activation (27), while Rojas-Lopez et al. found no significant involvement of NLRP11 in this process (26). Our results confirm a role for NLRP11 in activation of the non-canonical inflammasome and demonstrate that the NACHT and/or LRR domains are required for NLRP3 activation but that NLRP3 activation does not strictly depend on the NLRP11 pyrin domain (Fig. 1C-D). With respect to a role for NLRP11 in activation of the NLRP3 inflammasome specifically, we speculate that the difference in the results reported herein compared with those reported in Rojas-Lopez et al. is a result of some genomic instability of the cell lineage in Rojas-Lopez et al. due to the integration of *Cas9*. The new constructs described in the present study were generated by transfection of Cas9 protein together with guide RNAs (Materials and Methods). The constitutive presence of *Cas9* may have resulted in off-target effects that masked the role of NLRP11 specifically in the canonical NLRP3 inflammasome pathway. Our new results thus unite the findings in the field and establish a consensus that NLRP11 is required for activation of both the non-canonical and the canonical NLRP3 inflammasome.

Although the mechanisms by which NLRP11 promotes NLRP3 activation remain unclear, we speculate that NLRP11 may function as a helper NLR, as has been described for both mammalian and plant NLRs. In some cases, as NLR that has evolved to recognize the pathogen-associated molecular pattern interacts with a second NLR that executes immune signaling (37–41). In other cases, two NLRs that interact biochemically function more efficiently as a hetero-complex (37, 42, 43). The observation that NLRP11 is present in human macrophages at very low levels (26) suggests that NLRP11 may act upstream of NLRP3, which is abundant, rather than forming a hetero-complex. However, future investigations are required to clarify the mechanisms of NLRP11 function in NLRP3 activation.

The murine homologue of human caspases-4 and -5, caspase-11, is required for the detection of cytosolic LPS in mice. *Caspase-11*^-/-^ mice are more resistant to death from *Escherichia coli*-induced or LPS-induced septic shock (22, 24). As described above, in humans, caspase-4 and/or caspase-5 (25, 44, 45) and NLRP11 (26, 27), which is absent in mice, are required for recognition of cytosolic LPS. Indirect evidence suggests that lipoteichoic acid (LTA) of gram-positive bacteria can also be detected by caspase-4 (46, 47). Here, we demonstrate for the first time that NLRP11 and the non-canonical caspase-4 inflammasome are required for detection of intracellular *M. tuberculosis* (Fig. 2, Fig. 4, and Fig. 5). Our results indicate that, for infection with *M. tuberculosis* or *M. kansasii*, sensing of the infection by the non-canonical pathway is essential for host inflammasome responses, since caspase-4/5 are required for IL-1β release (Fig. 5). Interestingly, a 2010 shRNA screen of NOD proteins required for IL-1β release from *M. tuberculosis*-infected THP-1 cells identified *NLRP11* as a hit (48).

*M. tuberculosis* contains neither LPS nor LTA, and so what could the ligand be for caspase-4 and/or NLRP11? Potential ligands include the lipoarabinomannans (LAM), which are complex glycolipids similar to LPS and, like lipid A of LPS, are anchored in the inner membrane via a lipid anchor (33). The sugar composition of LAMs differs among mycobacterial species (49), which could explain why *M. tuberculosis* and *M. kansasii* but not *M. smegmatis* induce non-canonical inflammasome activation (Fig. 4). Future investigation of the role of LAM in inflammasome activation is needed. A second possible group of mycobacterial ligands for caspase-4 and/or NLRP11 are the Pro-Glu motif-containing and Pro-Pro-Glu motif-containing (PE/PPE) domain containing family of proteins, which are abundant in mycobacterial species such as *M. tuberculosis* and *M. kansasii* but absent in *M. smegmatis* (50, 51). However, the PPE protein PPE13 interacts with the NACHT or LRR domains of NLRP3, leading to its activation (52), whereas our data indicate that direct activation of NLRP3 is unlikely the mechanism of action for *M. tuberculosis* and *M. kansasii*, since we found an almost complete absence of IL-1β secretion in caspase-4/5 deficient THP-1 cells (Fig. 5). In conclusion, our results expand the range of pathogens recognized by NLRP11 and caspase-4 to now include acid-fast bacteria and underscore the versatility of these innate immune components in pathogen detection.

## MATERIALS AND METHODS

### Bacterial strains and growth conditions

*M. tuberculosis* (H37Rv), *M. kansasii* (ATCC1278), and *M. smegmatis* (mc2155) strains were grown in liquid Middlebrook 7H9 medium supplemented with 10% oleic acid-albumin-dextrose-catalase (OADC) growth supplement (B12351), 0.2% glycerol, and 0.05% Tween 80. *S. flexneri* wild-type serotype 2a strain 2457T was used in this study (53). *S. flexneri* was grown in tryptic soy broth at 37°C with aeration.

### Generation of CRISPR deletions in NLRP11

We previously created a deletion of the coding sequences of the NACHT and LRR domains in the *NLRP11* gene of THP-1 monocytes using a lentivirus-based system to deliver guide RNAs into cells in which *Cas9* had previously been stably integrated, such that the generated *NLRP11* knockout carried an integrated copy of *Cas9* gene (26). In the current study, to generate knockout lineages that lack chronic *Cas9*, chemically modified sgRNAs (Synthego) were electroporated as ribonucleoprotein (RNP) complexes into freshly cultured THP-1 monocytes using the Neon Transfection System (Invitrogen, MPK5000) following the manufacturer’s instructions. Specifically, to create the editing RNP complexes, 100 pmol of each sgRNA (Fig. S1E) were mixed with 25 pmol of SpCas9 2NLS nuclease (Synthego) (4:1 ratio) in Resuspension Buffer R (Invitrogen, MPK1069B) and incubated for 15 min at room temperature. THP-1 monocytes were washed in PBS and resuspended in Buffer R. The resulting RNP complexes were electroporated by pulsing twice 1x10^6^ cells in the 10 µL Neon Tip at 1700 V and 2 ms pulse width. Electroporated cells were immediately resuspended in 2.5 mL complete growth medium and placed into a 6-well plate. Cells were incubated for 72 hours at 37°C, after which 1.5 mL was transferred to a 75 cm^2^ culture flask, and 1 mL was used to purify genomic DNA using DNeasy Blood and Tissue kit (Qiagen). Following PCR and sequencing, editing efficiency was determined using ICE Analyzer (Synthego). The remaining cells were limiting-diluted and expanded as single clones in a 96-well dish. Using sequencing, isolated single clones were validated for homozygous out-of-frame deletions (Fig. S1).

### Mammalian cell culture

WT human-derived THP-1 monocytes (American Type Culture Collection, TIB-202) were obtained by ATCC. Caspase-4 and -5 deficient THP-1 cells were generated as described (54, 55) and kindly provided by Dr. C.Y. Taabazuing (University of Pennsylvania). Cells were maintained in RPMI 1640 medium (Gibco, 11875119) supplemented with 10% (v/v) fetal bovine serum (heat-inactivated, sterile-filtered, R&D systems) and 10 mM HEPES (Gibco, 15630-080). Cells were grown in 5% CO_2_ in a humidified incubator at 37°C. For use in experiments, cells were seeded in 48-well plates at 2×10^5^ cells per well or in 96-well plates at 5×10^4^ cells per well. For all non-mycobacterial experiments, to induce differentiation into macrophages, cells were incubated with 100 ng/mL phorbol 12-myristate 13-acetate (PMA) for 24 h, after which they were rested in media without PMA overnight prior to use in experiments. For mycobacterial infections, cells were differentiated into macrophages in 50 ng/mL PMA for 48 h, after which they were rested in media without PMA for 48 h prior to use in experiments.

### Cell treatments

In experiments with nigericin treatment, PMA-differentiated THP-1 macrophages were primed with 500 ng/mL ultrapure LPS from *Escherichia coli* O111:B4 (InvivoGen, tlrl-3pelps) for 4 hours followed by treatment with 10 µM nigericin (InvivoGen, tlrl-nig) for 90 min. In experiments involving nigericin treatment or *S. flexneri* infection, 2 µM MCC950 (NLRP3 inhibitor, InvivoGen, inh-mcc) was added where appropriate one hour prior to addition of nigericin or infection and maintained in the medium for the duration of the experiment. In cells infected with *M. tuberculosis*, MCC950 was added at 4 h of infection and maintained in the medium for the duration of the experiment.

### Bacterial infections

For infections with mycobacterial strains, bacteria grown to an exponential phase were centrifuged at 2000 g for 7 min. Bacterial pellets were resuspended in 1X phosphate-buffered saline (PBS) containing 0.05% Tween 80 and centrifuged at 80 g for 3 min to pellet clumped bacteria. The supernatants (single cell bacterial suspensions) were used to determine the multiplicity of infection (MOI) by optical density. PMA differentiated THP-1 macrophages were infected at the MOI of 10 and incubated for 2 h (*M. smegmatis*) or for 4 h (*M. tuberculosis* and *M. kansasii*) in 5% CO_2_ in a humidified incubator at 37°C. After this period of phagocytosis, infected macrophages were gently washed twice with prewarmed 1X PBS. Gentamicin, to kill extracellular bacteria, was added at 100 µg/mL in complete media containing RPMI1640 supplemented with 5% human AB serum. Samples for LDH release assay and IL-1β ELISA were collected at the indicated times.

For infections with *S. flexneri*, PMA-differentiated macrophages were infected at an MOI of 10 with bacteria grown to exponential phase after back-dilution from an overnight culture. Bacteria were centrifuged onto macrophages at 800 g for 10 min at room temperature, followed by incubation in 5% CO_2_ in a humidified incubator at 37°C. After 30 min, cells were washed twice with Hanks’ Balanced Salt Solution (HBSS, Gibco, 14025-092), and gentamicin was added to 25 µg/mL in RPMI 1640 medium supplemented with 10 mM HEPES without serum. Samples for LDH release assay, cytokine ELISAs, and western immunoblot were collected at 2 h 40 min. of infection.

### LDH cytotoxicity assay

Macrophage cell death was assessed by release of lactate dehydrogenase (LDH) due to loss of cellular membrane integrity. To measure LDH activity, 50 μL of cell culture supernatants from a 96-well plate were collected at indicated times and a CytoTox 96® Non-Radioactive Cytotoxicity Assay (experiments with mycobacteria; Promega, G1780) or a CyQUANT™ LDH Cytotoxicity Assay (experiments with *S. flexneri* and nigericin; ThermoFisher Scientific, C20301) was used following the manufacturer’s instructions.

### Cytokine ELISAs

Cell culture supernatants harvested from mycobacteria-infected cells were assayed for levels of secreted IL-1β using Human Total IL-1β DuoSet ELISA kit (biotechne, DY201) following the manufacturer’s instructions. Cell culture supernatants harvested from cells infected with *S. flexneri* or treated with nigericin were assayed for levels of secreted cytokines using Human Total IL-18 DuoSet ELISA kit (biotechne, DY318-05) and ELISA MAX™ Deluxe Set Human IL-1β kit (BioLegend, 437004) following the manufacturers’ instructions.

### Protein extraction and western immunoblot analysis

For detection of released caspases, 100 μL of cell culture supernatants from a 48-well plate were collected at the indicated times and mixed with 35 μL of 4x Laemmli buffer [250 mM Tris (pH 6.8), 8% SDS, 50% glycerol, 0,04% bromophenol blue] supplemented with cOmplete EDTA-free protease inhibitor (Roche, 11873580001). For detection of cell-associated caspases and full-length and cleaved GSDMD, cells in the same plate were lysed directly in 100 μL of 4x Laemmli buffer supplemented with the same protease inhibitor. Following SDS– polyacrylamide gel electrophoresis (SDS-PAGE), proteins were transferred onto a nitrocellulose membrane.

Antibodies used for immunoblotting were as follows: caspase-1 (Abcam, ab207802) rabbit monoclonal antibody at 0.5 μg/mL (1:1000), caspase-4 (Santa Cruz, sc-56056) and GSDMD (Santa Cruz, sc-81868) mouse monoclonal antibodies at 0.5 μg/mL (1:200). Secondary antibodies were goat anti-rabbit immunoglobulin G (IgG) or goat anti-mouse IgG, each conjugated to horseradish peroxidase (HRP) (Jackson ImmunoResearch). Antibodies specific for β-actin were HRP-conjugated (Sigma, A3854). Immunoreactive bands were visualized by chemiluminescence with SuperSignal West Pico PLUS or SuperSignal West Femto Maximum Sensitivity substrates (Thermo Fisher Scientific, 34580 and 34096).

### Statistical analysis

GraphPad Prism software was used for graphing data and statistical analyses. Statistical significance was determined using the ordinary two-way ANOVA with Tukey’s multiple comparisons test, with a single pooled variance. For Fig. 4, statistical significance was determined using a paired parametric t-test. Data were graphed so that each data point represents the mean of technical triplicate wells for one experiment, and all results are data from at least three independent biological replicates, unless noted otherwise. Differences were considered statistically significant if the *P* value was <0.05.

## ACKNOWLEDGMENTS

This work was supported by NIH R01 AI173030 (to M.B.G.), NIH R01 AI081724 (to M.B.G.), and NIH R01 AI147630 (to V.B.), NIH T32 AI007061 (to M.S.), and an MGH ECOR Fund for Medical Discovery Fundamental Research Fellowship Award (to M.L.G.M.).

## Notes

### Competing Interest Statement

The authors have declared no competing interest.

### Summary of Updates

The manuscript has been revised for clarity. A summary figure has been added to better visualize the findings of the study.

