## Supplemental materials for "NLRP11 is required for canonical NLRP3 and non-canonical inflammasome activation during human macrophage infection with mycobacteria"

### SUPPLEMENTAL FIGURES

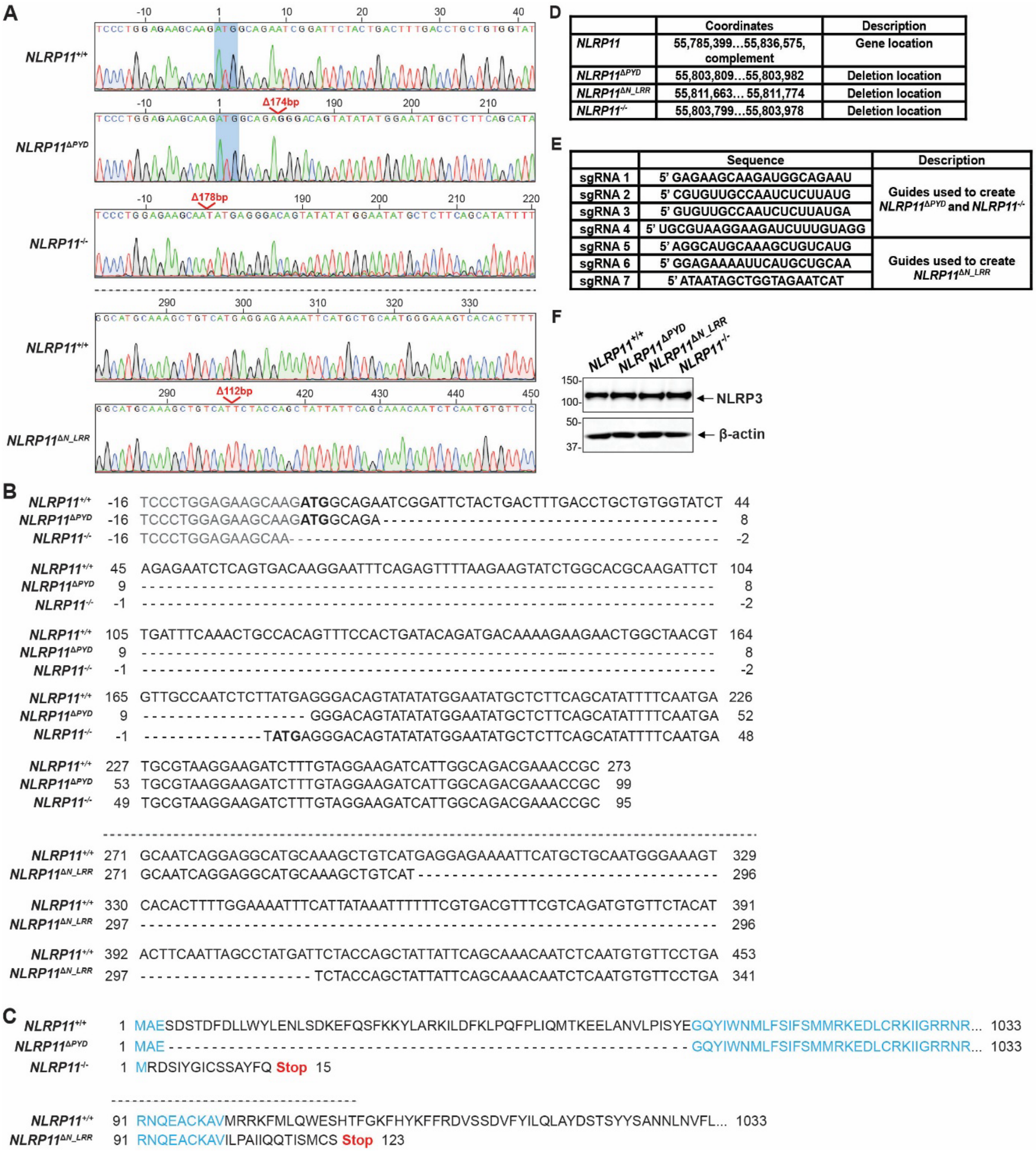

**FIG S1** Generation and validation of NLRP11 deletions. (A) Sequencing analysis following PCR amplification of appropriate portions of the *NLRP11* gene in clonally expanded indicated THP-1 monocytes. Start codon is marked by blue shadowing. Generated deletions are indicated in red. Relative position to the start codon is indicated. The data show that the indicated deletions are homozygous. (B) Alignment of the DNA sequences of the generated *NLRP11* deletion mutants. Start codons are shown in bold. Intron sequences are shown in gray. Relative position to the start codon is indicated. (C) The predicted amino acid sequence and protein length of the resulting NLRP11 proteins are shown. Sequences identical between the wt and mutant proteins are marked in blue. Premature stop codons are indicated. (D) Genomic location (chromosome 19, GRCh38.p14, Primary Assembly) of the *NLRP11* gene (Gene ID: 204809; updated on November 27, 2024. NCBI Reference Sequence: NC_000019.10) and of the generated deletions. (E) Sequences of the sgRNAs used to generate the deletions in the NLRP11 gene. (F) NLRP3 levels in whole cell lysates of indicated macrophages. Western blot. Molecular weight (MW) markers are in kilodaltons.

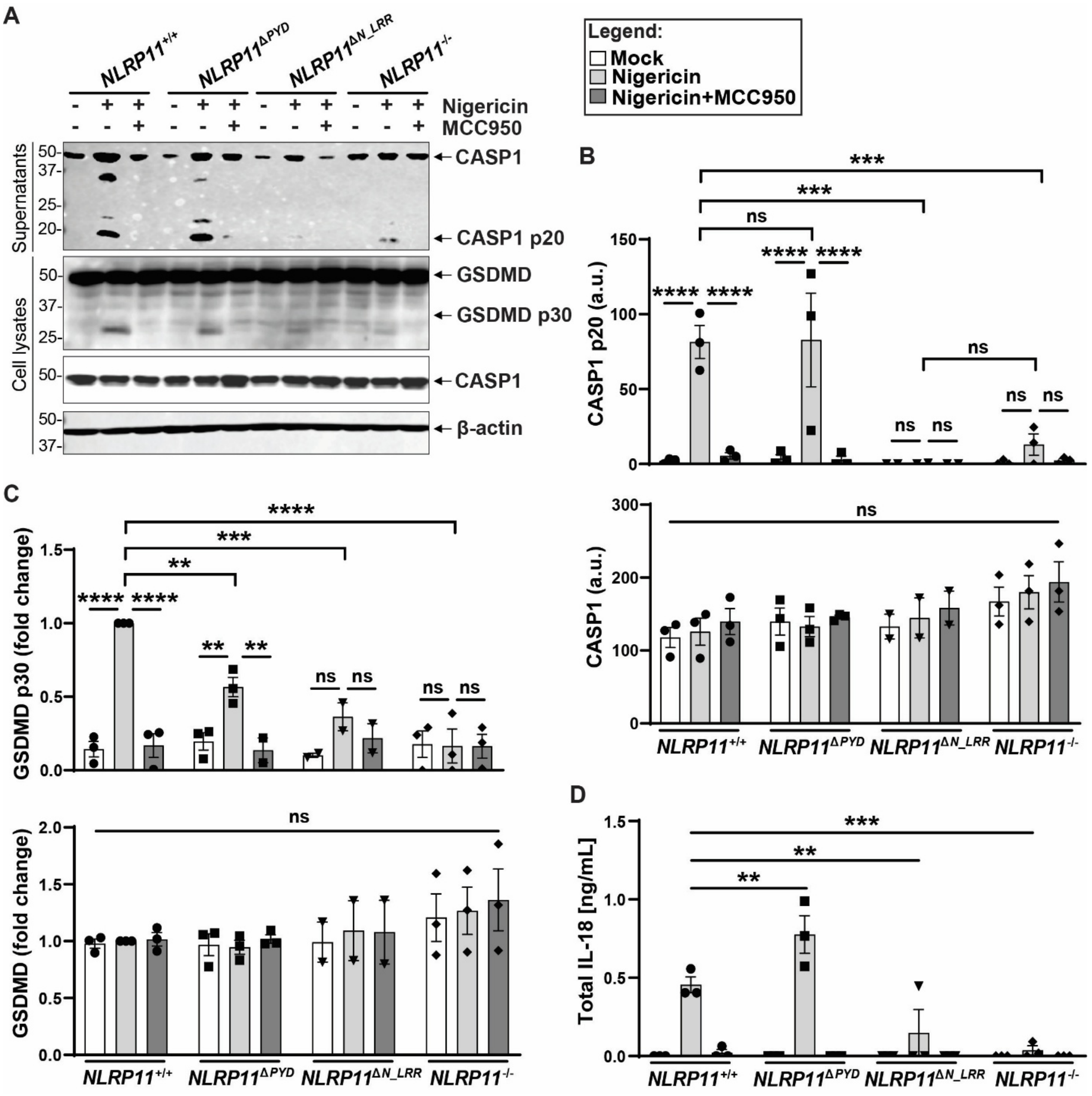


**FIG S2** Supplementary data to Fig. 1. (A) Cleaved caspase-1 (p20) released into the cell culture supernatants, and cleaved GSDMD (p30) in cell lysates in indicated PBS- or MCC950-pretreated macrophages at 90 min of nigericin treatment. Representative western blots. (B-C). Quantification (band densitometry) of immunoblots in panel A. Cleaved caspase-1 and GSDMD (top) were normalized to the full-length proteins, and full-length caspase-1 and GSDMD (bottom) were normalized to β-actin. Levels of caspase-1 are expressed in arbitrary units (a.u.) (B), whereas levels of GSDMD are expressed as fold change compared to nigericin-treated NLRP11$^{+/+}$ macrophages. (D) IL-18 released into the cell culture media from macrophages treated as in panel A, measured by ELISA at 90 min of nigericin treatment. ns, not significant; **, $P < 0.01$; ***, $P < 0.001$; ****, $P < 0.0001$; by two-way ANOVA. Shown are independent biological replicates (B-D). Error bars represent the standard error of mean.

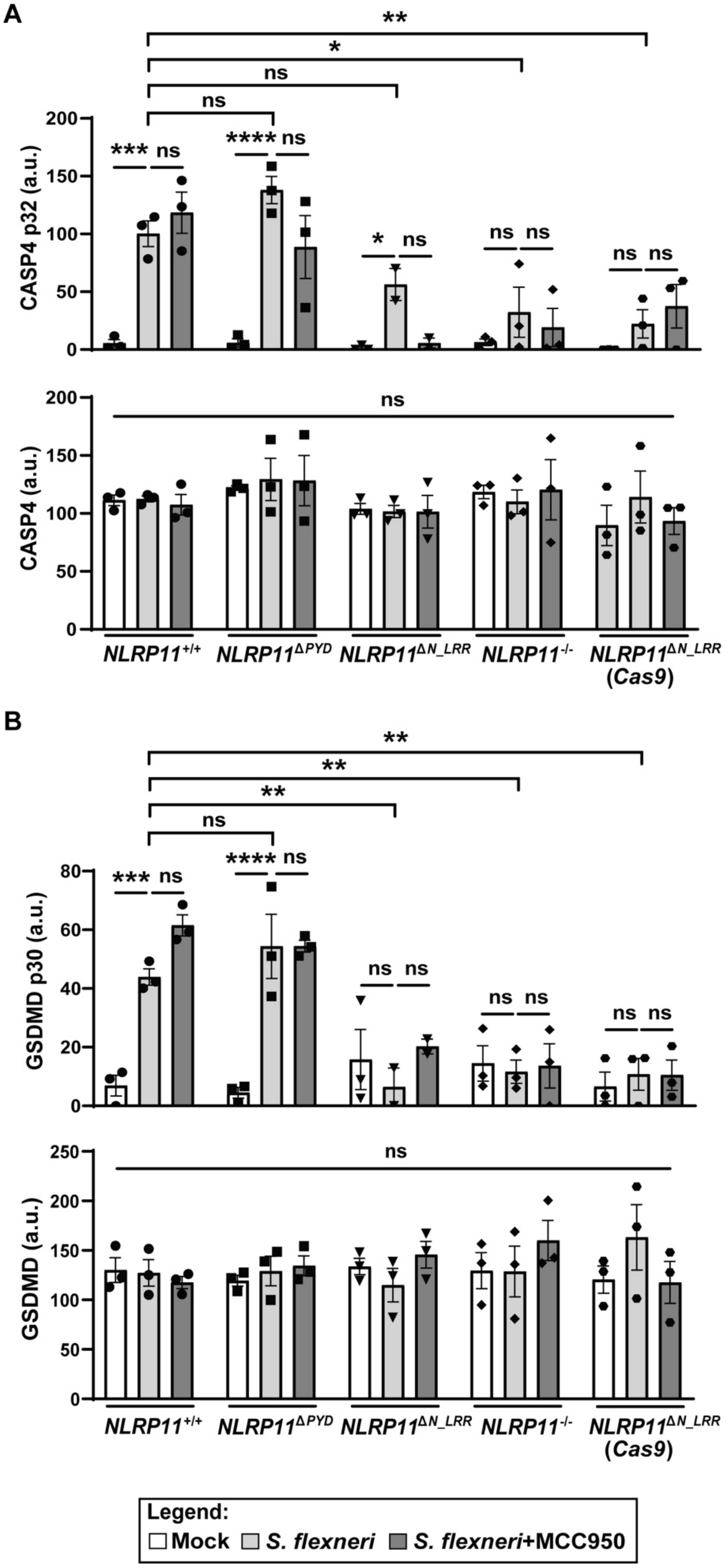
A
CASP4 p32 (a.u.)
CASP4 (a.u.)
NLRP11+/+
NLRP11ΔPYD
NLRP11ΔN_LRR
NLRP11-/-
NLRP11ΔN_LRR (Cas9)
B
GSDMD p30 (a.u.)
GSDMD (a.u.)
Legend:
Mock
S. flexneri
S. flexneri+MCC950

**FIG S3** Supplementary data to Fig. 3. (A-B) Quantification (band densitometry) of western immunoblots from Fig. 3C, measuring release of cleaved caspase-4 into cell culture supernatants (A), and cleaved GSDMD in cell lysates (B) in indicated PBS- or MCC950-pretreated macrophages at 3 h of infection with *S. flexneri* (MOI 10). Cleaved caspase-4 and GSDMD (top) were normalized to the full-length proteins, and full-length caspase-4 and GSDMD (bottom) were normalized to β-actin, expressed in arbitrary units (a.u.). ns, not significant; *, $P < 0.05$; **, $P < 0.01$; by two-way ANOVA. Shown are independent biological replicates. Error bars represent the standard error of mean.

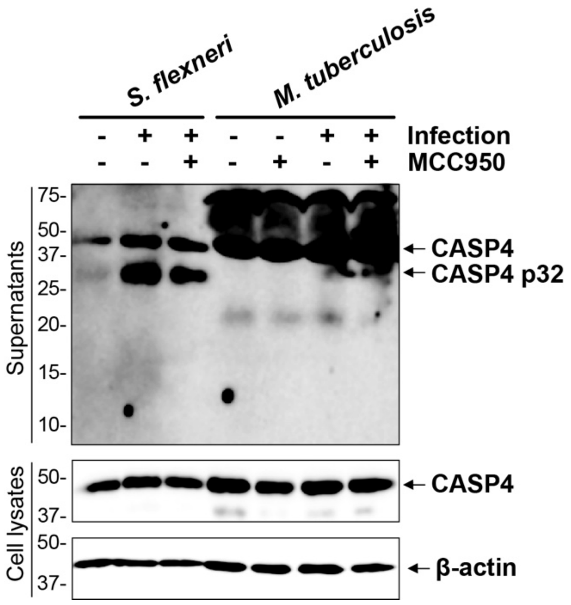


**FIG S4** Supplementary data to Fig. 5. Cleaved caspase-4 (p32) released into the cell culture supernatants and full-length caspase-4 in cell lysates in *M. tuberculosis*-infected (MOI 10, 8h of infection) or *S. flexneri*-infected (MOI 10, 3h of infection) WT THP-1 macrophages, pretreated or not with MCC950. Representative western blots.
